# Critical Pressure of Intramural Delamination in Aortic Dissection

**DOI:** 10.1101/2021.09.12.459981

**Authors:** Ehsan Ban, Cristina Cavinato, Jay D. Humphrey

**Affiliations:** Department of Biomedical Engineering, Yale University, New Haven, CT, USA

**Keywords:** delamination, phase field, intra-lamellar strength, tearing pressure

## Abstract

Computational models of aortic dissection can examine mechanisms by which this potentially lethal condition develops and propagates. We present results from phase-field finite element simulations that are motivated by a classical but seldom repeated experiment. Initial simulations agreed qualitatively and quantitatively with data, yet because of the complexity of the problem it was difficult to discern trends. Simplified analytical models were used to gain further insight. Together, simplified and phase-field models reveal power-law-based relationships between the pressure that initiates an intramural tear and key geometric and mechanical factors – insult surface area, wall stiffness, and tearing energy. The degree of axial stretch and luminal pressure similarly influence the pressure of tearing, which was ∼88 kPa for healthy and diseased human aortas having sub-millimeter-sized initial insults, but lower for larger tear sizes. Finally, simulations show that the direction a tear propagates is influenced by focal regions of weakening or strengthening, which can drive the tear towards the lumen (dissection) or adventitia (rupture). Additional data on human aortas having different predisposing disease conditions will be needed to extend these results further, but the present findings show that physiologic pressures can propagate initial medial defects into delaminations that can serve as precursors to dissection.

## Introduction

Aortic dissection is a life-threatening event that is increasingly responsible for significant morbidity and mortality. Despite considerable clinical experience, histo-pathological characterizations, *in vitro* biomechanical findings, and computational simulations,^15,34,40^ the precise mechanisms by which the aorta dissects remain unknown. It is axiomatic, however, that dissections arise when mechanical stresses exceed material strength, hence emphasizing the importance of performing and interpreting mechanical tests on the aorta. Not surprisingly, many different types of experiments have been reported, including uniaxial tests to failure, inplane shearing tests, so-called peeling tests on strips of tissue, and pressurization to failure of cylindrical segments. Peeling and in-plane shearing tests provide particularly important information on intra-lamellar adhesion strength,^28,35^ but these tests impose non-physiological loading conditions on a non-physiological geometry. Among others, Margot Roach and colleagues presented results from a clever experiment that preserved the native cylindrical geometry and *in vivo* relevant biaxial loading conditions while examining intra-lamellar adhesion – they used a 25-gauge needle to inject pressurized ink into the media of excised but intact pressurized segments of healthy porcine aorta.^30^ They found that the mean pressure of tearing, defined as the difference between the imposed intra-lamellar pressure and the prescribed luminal pressure, was on the order of 547 mmHg (∼73 kPa) while the mean pressure at which the delamination propagated was 54 mmHg (∼7.2 kPa), independent of the depth within the media at which the tip of the needle rested. Because of the complexity of the mechanics associated with this experiment, their findings were discussed primarily in qualitative terms. Given the potential importance of these findings, there was a need to revisit these data and to seek increased understanding, particularly for human aortas.

In this paper, we present finite element-based simulations of experiments such as those reported by Roach and colleagues to gain new insight into the many factors that dictate the propagation of a tear within the aortic wall. Notwithstanding the advantages of our phase-field approach, the complexity of this three-dimensional boundary value problem yet rendered it initially difficult to gain broad intuition. Hence, we also solved multiple highly idealized boundary value problems (using adherent beams, plates, and membranes) that reflect different aspects of the complex experiment of interest, thus increasing insight into both the mechanisms of failure and the different factors – degree of the initiating insult, wall stiffness, and fracture properties – that drive delamination of the aortic media. Finally, we examined computationally the potential impact of focal inhomogeneities of wall properties on the direction of the propagation. Collectively, these findings provide new clues regarding both dissection (i.e., tears that turn inward toward the lumen) and transmural ruptures (i.e., dissections that turn outward toward the adventitia).

## Methods

### Phase-field finite element model of intra-lamellar tearing

Two- and three-dimensional finite element models were used to simulate the *in vitro* injection of ink into the medial layer of cylindrical segments of excised aorta subjected to axial extension and luminal pressurization. Specifically, we used a phase-field finite element framework that was validated in a previous study of ink injection into cut-open, traction-free flat segments of aorta^4^. This phase-field model of tearing is based on a global minimization of energy,^11^ with tearing defined by a phase-field *ϕ* ∈ [0,1) where *ϕ* = 0 and 1 correspond to intact and fully damaged tissue, respectively.

The finite element mesh in the three-dimensional model consisted of a cylindrical domain divided into ∼500,000 tetrahedral elements, generated using an in-house script. The displacement field ***u*** was defined over the domain, and isochoric deformations were enforced using a Lagrange multiplier field *p*. Only a quarter of the three-dimensional vessel geometry was modeled, with zero axial displacements prescribed at one end of the cylindrical domain, as in the biaxial tests on intact segments, and symmetric boundary conditions on the other end. These symmetries restrict the propagation of possible tears to symmetric paths. Residual stresses were accounted for naturally by prescribing constituent-specific deposition stretches within a four-fiber family pseudoelastic constitutive relation and ensuring equilibrium in the traction-free configuration.^33^ These deposition stretches were 1.2 for elastin and 1.08 for each family of collagen fibers. The two-dimensional model was generally similar, but focused on a single cross-section. We used ∼37,000 triangular elements in these simulations. In both cases, luminal pressure was applied as a traction boundary condition over the luminal surface (Fig. 1(a), (b)). The applied pressure at the current increment of loading remained normal to the deformed luminal surface, evaluated at the previous increment. To initiate a computation, axial stretch was increased gradually from 1 to the in vivo value, then the luminal pressure was increased gradually from 0 to 130 mmHg to represent the arterial pressure prescribed experimentally.^30^ This initialized model included a small damaged volume that corresponded to a torn surface area created by the insertion of the needle (Fig. S1 in Supplemental Information). Injection of ink into the arterial wall was modeled by incrementally increasing the volume of the ink, *V*_injection_, using a global Lagrange multiplier, *m*.^4^ Tears emerged as concentrated regions of damage surrounded by elastic material, similar to previous analyses of rubber-like elastomers and arterial tissue.^20^

**Figure 1.**
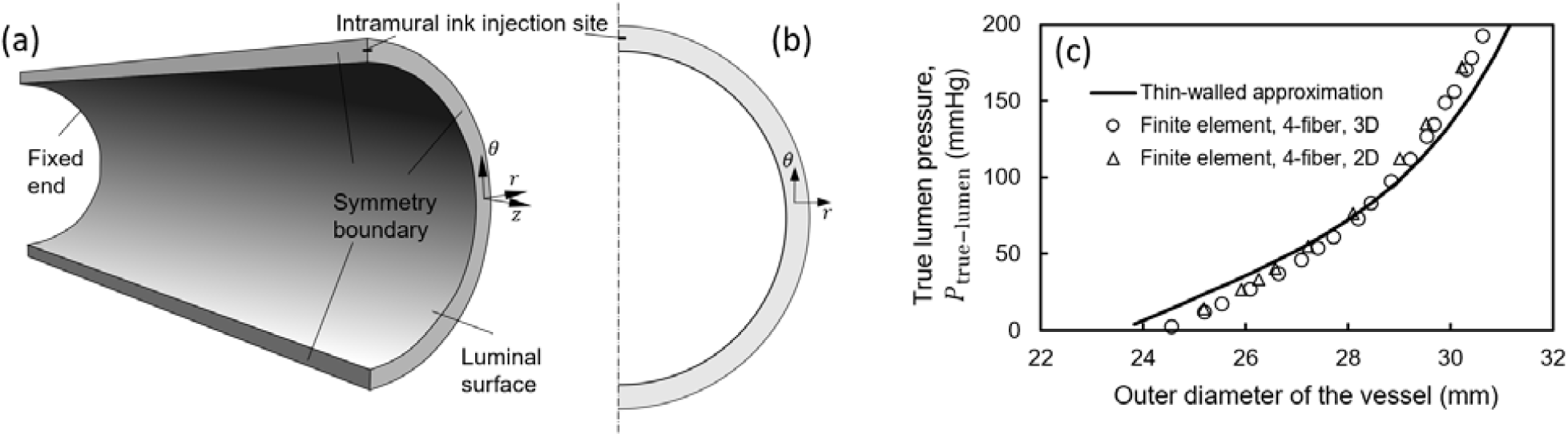
Model setup for propagating an intramural tear in a biaxially loaded human aorta by an intramural injection of ink. (a) Three-dimensional model showing an arterial section, modeled using symmetric boundary conditions. Zero axial displacements were prescribed at the two ends of the initially axially extended and pressurized vessel, thus resembling a biaxial test setup, and symmetri displacement boundary conditions were prescribed at the other cross-sectional boundary surfaces. The volume of the damaged tissue was prescribed in increments beginning with the injection site (Fig. S1), which models the incision made by the needle. (b) Two-dimensional model of injection into the aortic wall. Symmetric displacement boundary conditions were employed in this model as well. (c) Verification of the implementation of the two- and three-dimensional Fung-type four-fiber family constitutive relations in the finite element code by comparison with an analytical thin-walled approximation of wall deformation during biaxial loading of a healthy human aorta, shown as the pressure of the true lumen versus the outer diameter of the vessel at various pressures.

At the *j’* th increment of injection volume, the fields ***u***_*j*_, *p*_*j*_, *ϕ*_*j*_ and *m*_*j*_ are determined by an alternate minimization of the total energy

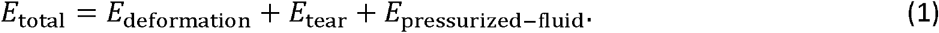

*E*_deformation_ denotes the energy due to deformation of the aortic wall, consisting of three contributions: the strain energy, a term that enforces local isochoric deformations of the wall, and a regularization term (Eqs. S1-S2 in Supplemental Information). *E*_tear_ is the energy of tearing, that is, the characteristic energy of tearing, *G*_*c*_, multiplied by a function of *ϕ* that represents the damaged area (Eq. S3 in Supplemental Information). A value of 52 J/m^2^ was used for *G*_*c*_, which lies within the range of values previously reported by others.^28,31,34^ We incrementally prescribed the volume of injection by the minimization of *E*_pressurized – fluid,_

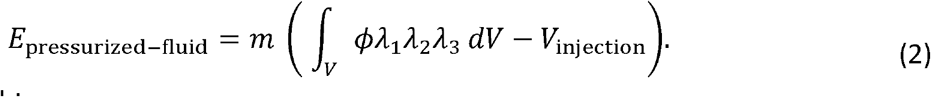

Finally, we sought

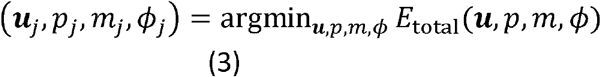

by iterative minimization, finding solutions of the equations 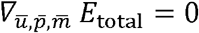 and 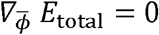, where 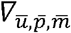 denotes a directional derivative with respect to ***u***, *p*, and *m* in the direction of their respective test functions.^4^ Similarly, 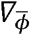 denotes the directional derivative with respect to *ϕ* in the direction of the test function for *ϕ*. That 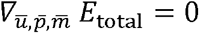 and 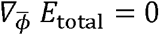 constitutes a weak form solution of the problem. The solution for the progressive damage was constrained between *ϕ* at the previous increment of injected volume and its maximum value of 1, thus enforcing irreversible damage via tearing. We implemented the Bubnov-Galerkin method^22^ within the FEniCS finite element framework.^2^ Mesh convergence was confirmed in preliminary simulations (Fig. S2 in Supplemental Information) for these fully (materially and geometrically) nonlinear problems.

### Fung-type constitutive model of the aortic wall with four families of fibers

In particular, the native aortic wall was modeled using an incompressible hyperelastic constitutive relation that accounts for overall structural contributions of elastin, collagen, and smooth muscle. A neo-Hookean contribution to the strain energy models the deformation of the elastin-dominated amorphous material. Collagen fibers and passive smooth muscle were modeled as Fung-type exponentially strain-stiffening materials.^33^ The constitutive model is defined by the strain energy density function

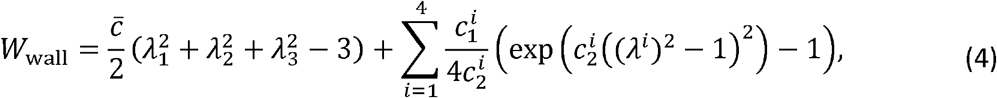

where 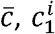, and 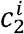 are material constants. The principal stretch ratios are denoted by *λ*_1_, *λ*_2_, and *λ*_3_, and the stretch ratios in the four main structural directions *i* are denoted by *λ*^*i*^, with *i* = 1,2,3, and 4 corresponding to axial, circumferential, and two symmetric diagonal directions, respectively. The diagonal directions lie within the axial-circumferential plane and are oriented at angles ± *α* with respect to the axial direction.

To increase clinical relevancy, a first set of illustrative material parameters was obtained from literature data for a healthy human descending thoracic aorta^17^ (Fig. 1(c)), as modeled previously,^33^ noting that porcine (studied by Roach and colleagues^30^) and young human (considered herein) aortas exhibit similar material behaviors. ^6^The inner radius and thickness of the healthy aorta were 11.75 mm and 1.75 mm, respectively.^33^ A second set of material parameters was used to model diseased aortic tissue based on available data,^20^ with radius and thickness of 15.0 mm and 2.5 mm. See Table S1 in Supplementary Information for specific values of these material parameters. Fig. S3 in Supplemental Information provides further information on the modeling of possible material inhomogeneities therein.

### Simplified analytical model

To develop intuition and gain qualitative understanding, we also developed simple extensions of previous analytical models of two interleaved tissue flaps and two fluid pools to study the pressure of tearing within a false lumen relative to the pressure of the true lumen. This study included the dependence of the pressure of tearing on the material properties. We previously used similar models^18^ to study delamination by injection of cut-open aortic tissue, that is, without a counter-acting true lumen pressure. These extended models included initially adherent linearly elastic structures (slender beams, circular plates, and circular membranes) that were separated by the action of the pressures applied to model the false lumen while including effects of true luminal pressures – see Appendix for key equations suggesting power-law relations and Supplemental Information (Figure S4 and Eqs. S4-S14) for additional details on solutions to such boundary value problems within a linear elastic framework. Again, we emphasize that these solutions were sought to gain qualitative understanding, not to model the actual experiment which is nonlinear geometrically (large strains) and materially.

### Experimental validation tests

To qualitatively confirm two novel computational findings (see Results), we repeated experiments of Roach and colleagues using needles of two different sizes, namely, 30- and 25-gauge. Eighteen descending thoracic aortas of similar length (56±3 mm) were harvested from healthy adult male Yorkshire pigs (22–27 kg), used in separate studies that did not affect the aortic mechanics or microstructure. Those studies were performed in accordance with protocols approved by the Institutional Animal Care and Use Committee of Yale University. As in the finite element models, the aortas were initially extended axially (to stretch ratios of 1.1 and 1.3) and pressurized to physiologic levels (130 mmHg), then injected locally with ink. Details on these experiments, including the physical setup (Fig. S5), are in Supplementary Information.

## Results

### Tears propagate suddenly after reaching a threshold injection pressure

The fully nonlinear three-dimensional phase-field model included a damaged area representative of the initial tear made by inserting a needle. As the volume of the injected ink increased, the damaged area initially grew larger and deformed the surrounding tissue without propagating the initial tear. Then, at a critical pressure, of fluid, *P*_tear_, the tear started to propagate, accompanied by a reduction in injection pressure (Fig. 2(a)). Upon further increase in the volume of injection, the tear propagated farther, increasing the torn area of the tissue (Fig. 2(b) and (c)). The tear propagated in both axial and circumferential directions, creating biconcave volumes. Although based on a geometry and material properties for a healthy human aorta, both the injection pressure versus volume curves and the morphology of the damaged vessels agreed well with those reported by Roach and colleagues for healthy porcine aortas.

**Figure 2.**
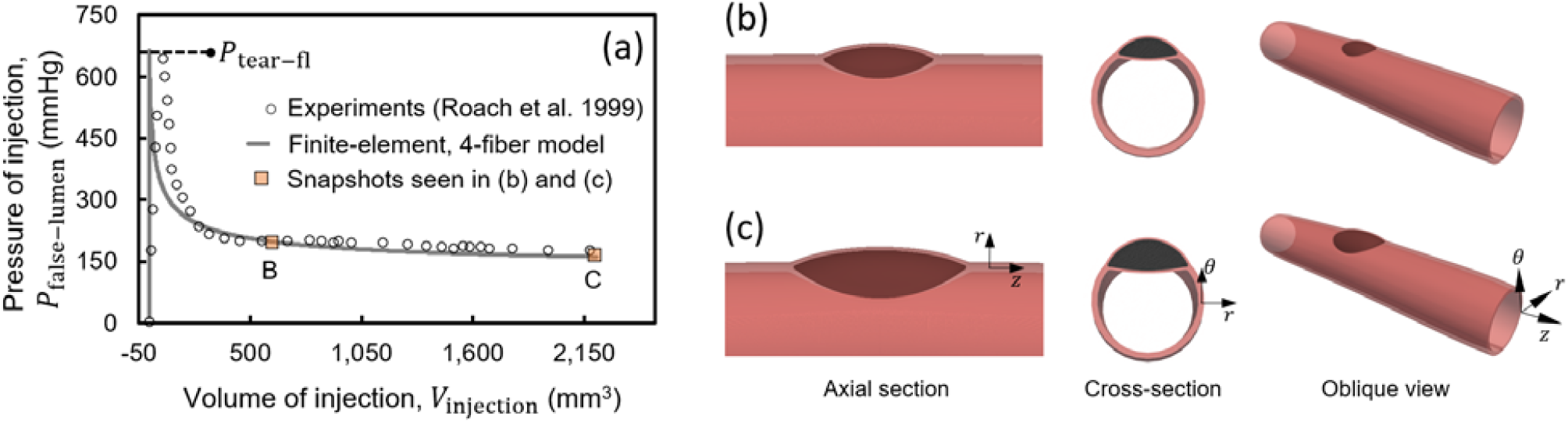
Finite element simulations of the intramural injection of ink within the media of a biaxially loaded vessel modeled by a Fung-type constitutive relation with four families of fibers for a healthy human aorta. (a) As ink was injected into the aortic wall, the pressure of injection initially increased rapidly while deforming the surrounding tissue. After reaching a critical value of pressure, *P*_tear_, the tear started to propagate axially and then circumferentially. Both the (a) injection pressure-volume curves and (b and c) the morphology of the injected volume agree qualitatively and quantitatively with those reported by Roach and colleagues.^30^

### Minimalistic analytical models suggest power-law relations

Together, the multiple simplified analytical models suggested that the pressure of tearing of the false lumen (*P*_tear-fl_) and the pressure of the true lumen (*P*_true lumen_) relate as (Appendix)

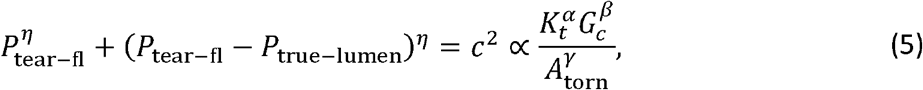

where *c* is related to the resistance of the tear to propagation, *K*_*t*_ is related to the tangent elastic stiffness, *G*_*c*_ is the characteristic energy of tearing, and *A*_torn_ is the torn area of the tissue (Fig. 3(a)–(g)). The exponents *α, β*, and *γ* take positive values but differ depending on the elastic structure in the model. In linear models with beams, plates, or highly tensed membranes, *η* = 2, thus at a constant resistance of the tear, *c*, Eq. (5) describes the relationship between *P*_tear-fl_ and *P*_true-lumen_ as the equation of an ellipse (curves in Fig. 3(h)). The analytical models with beams and plates model the bending deformation, more relevant to smaller torn areas, whereas models with membranes are more representative of the tension present in the deforming flaps at larger torn areas. The exponents were positive in all analytical and computational models.

**Figure 3.**
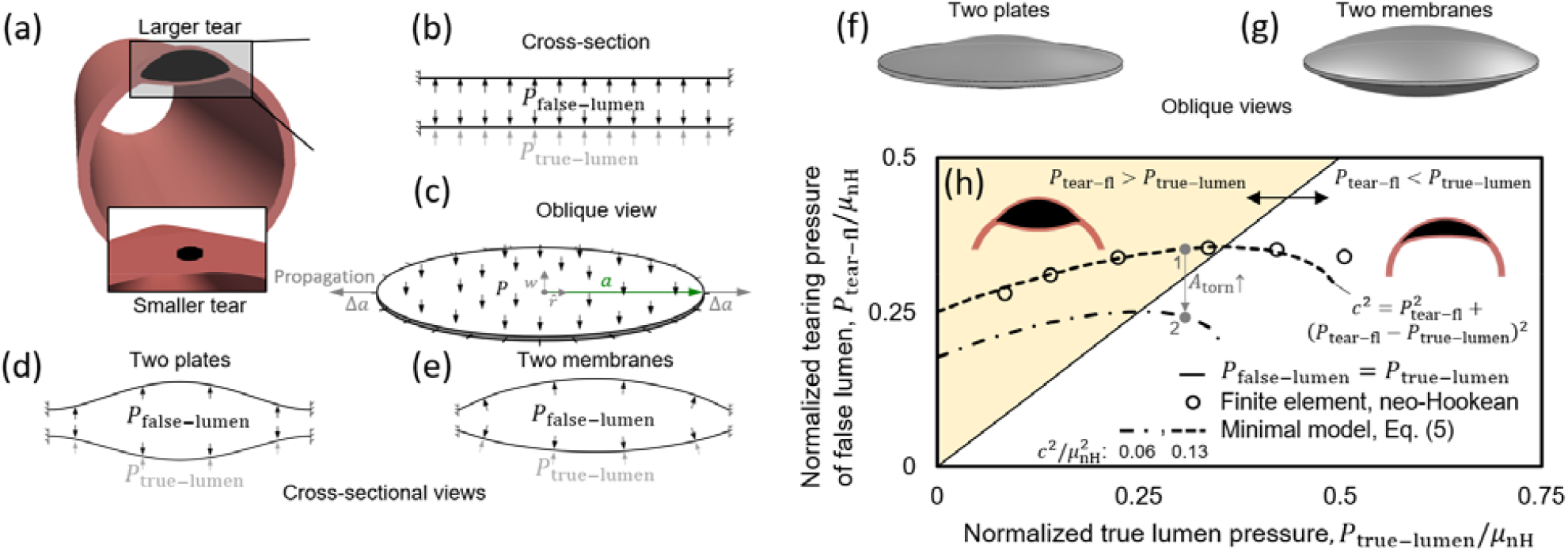
Schematic views of extensions of three minimalistic analytical models of the separation of adhered linear elastic structures with comparison to a nonlinear neo-Hookean finite element simulation. (a) Snapshot from a finite element model of intramural injection. A tear larger than the initial torn area in Fig. 2 is shown to highlight the tissue deformation and tearing via an exaggerated view. The initial insults due to the needle appear substantially smaller prior to the start of tear propagation. (b and c) Cross-sectional and oblique views of the loading of two portions of the wall by an intramural pressure, *P*_false-lumen_, and pressure in the true lumen, *P*_true-lumen_, in the minimalisti models. The models consist of initially adherent beams, circular plates, or circular membranes with zero displacements prescribed at their ends / periphery. (d and e) Cross sectional and (f and g) oblique schematic views of the deformation of the two (d and f) plates or (e and g) membranes. These highly simplified linear models suggest that the pressure of tearing at the false lumen relates to the pressure of the true lumen via an equation of an ellipse, 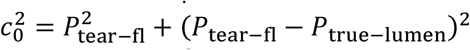. (h) Comparison of the results from a minimalistic model and a finite element simulation of a pressurized cylinder modeled as a neo-Hookean solid with an initial tear. The solid line indicates when pressures in the two lumens equal. Above the line, where *P*_false-lumen_ > *P*_true-lumen_, the larger pressure in the false lumen pushes the septum into the true lumen and leads to a biconcave shape (insets). Below the line, the pressure in the true lumen is larger, which leads to a crescent or meniscus shape of the false lumen. The trend of the slopes of the ellipses changes at their intersection with the *P*_false-lumen_ = *P*_true-lumen_ line.*P*_tear-fl_ increases with increasing *P*_true-lumen_ above the line; the reverse is observed below the line. The gray arrow shows the path going from point 1 to 2 with an increase in the original torn area of the wall. Interpretation of the model is limited to the regime of the injection experiments where the pressure of the false lumen is larger than that of the true lumen. These qualitative results motivated novel interpretations of simulations shown below for nonlinear finite element models of the aortic wall described by a Fung type four-fiber family constitutive model.

Detailed interpretation of these models is not recommended because of the use of linear elastic models. Nonetheless, qualitative insights emerge for intramural tearing of the wall by intramural injection. In the regime where *P*_tear-fl_ > *P*_true-lumen_ (shaded area in Fig. 3(h)), the false lumen has a biconcave morphology (Fig. 3(h), *inset*) and *P*_tear-fl_ increases with increasing *P*_true-lumen_. The needle injection experiments lie in this region. In contrast, if *P*_tear-fl_ < *P*_true-lumen_, the false lumen has a meniscus-like morphology (Fig. 3(h), *inset*) and *P*_tear-fl_ decreases with increasing *P*_true-lumen_. In the regime where *P*_tear-fl_ > *P*_true-lumen_ and its vicinity, the ellipse-shaped curve of the minimal model, Eq. (5), agrees reasonably well with results from finite element computations using a simplified nonlinear material description, a neo-Hookean model, thus suggesting utility of these minimalistic models. The main observations based on the analytical models are only qualitative, however, limited to the regime of the injection experiments where the pressure of the false lumen is larger than the pressure of the true lumen. In this regime, the critical pressure of tearing decreases with increasing size of the false lumen. Similarly, the critical pressure of tearing increases with increasing tissue stiffness and characteristic energy of tearing. Importantly, these minimalistic models suggest power-law relations that were thus investigated further based on full nonlinear finite element solutions described below.

### The pressure of tearing of the false lumen depends nonlinearly on the pressure of the true lumen, vessel stiffness, characteristic energy of tearing, and in vivo axial stretch

To probe changes of *P*_tear-fl_ with vessel properties, we again performed computations using the fully nonlinear phase-field finite element model with a four-fiber Fung-type constitutive relation for the healthy human aorta. First, *P*_tear-fl_ showed an increasing trend as the fixed pressure of the true lumen was increased incrementally from 0 to 130 mmHg. After that, it reached a plateau (Fig. 4(a)). Similarly, *P*_tear-fl_ decreased with decreases in the fixed value of axial stretch of the biaxially loaded vessel (Fig. 4(a), *inset*), which was confirmed qualitatively by the complementary experiments on porcine aortas that revealed a 1.5-fold change in *P*_tear-fl_ when testing at 1.1 and 1.3 axial stretch ratios (*p-value*=0.02). As expected, *P*_tear-fl_ increased withincreases in both the characteristic energy of tearing of the aortic wall and its native tangent stiffness (Fig. 4(b) and (c)). The exponents from Eq. (5) that relate tear resistance to *K*_*t*_ and *G*_*c*_ were found to be *α* = 1.0 and *β*= 1.5. Similar trends were observed in computations using constitutive relations for the diseased human aorta (Fig. 5).

**Figure 4.**
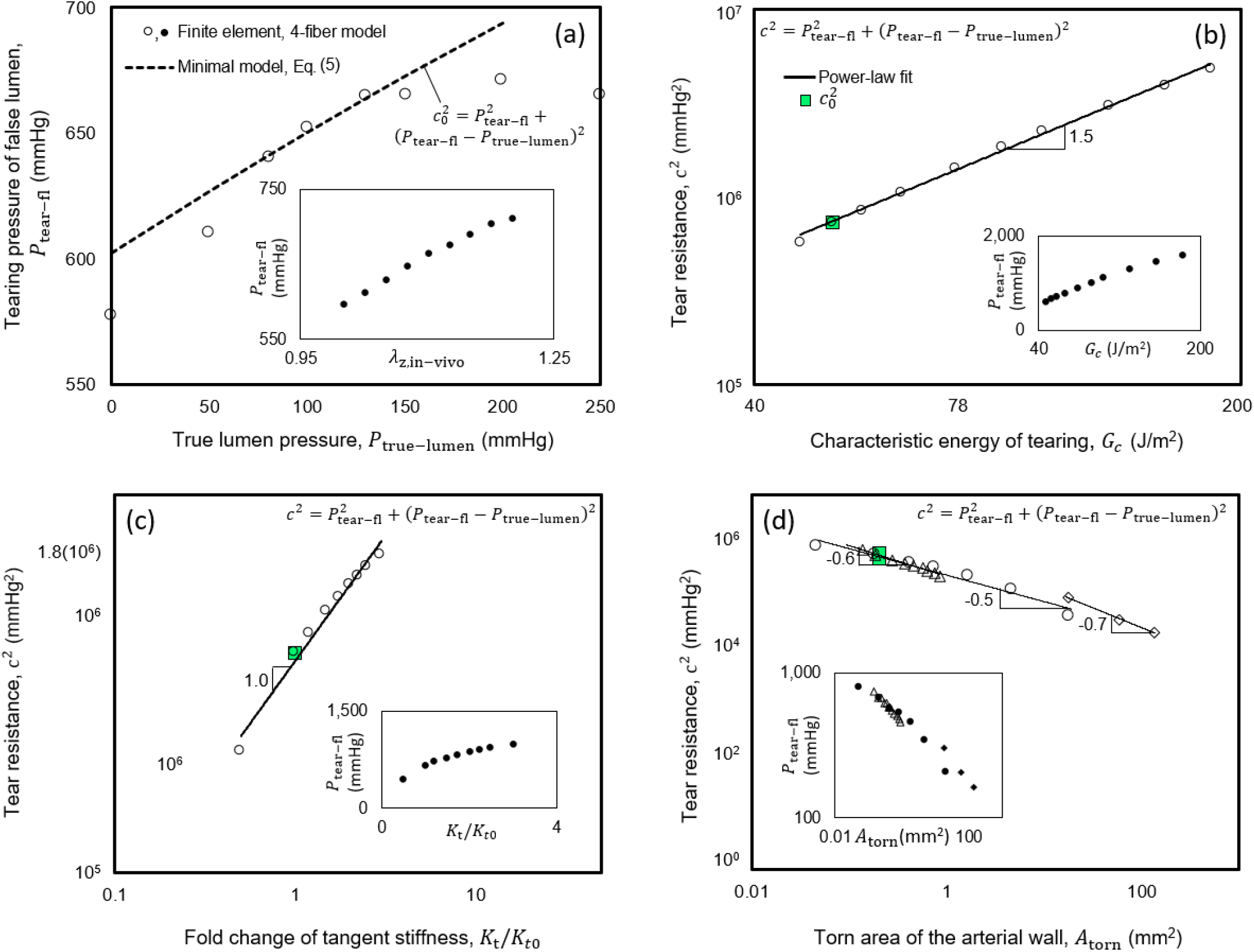
Dependencies of *P*_tear-fl_ on luminal pressure with tear properties assessed by the full nonlinear finite element model employing a Fung-type constitutive relation with four families of fibers for a healthy human aorta. (a) Variation of the pressure of tearing of the false lumen with the pressure of the true lumen. The dashed curve shows the fit to an equation of an ellipse, 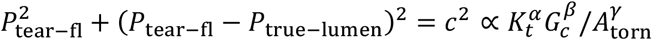, motivated by the minimalistic analytical models of linearly elastic materials. 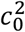 in this curve corresponds to the point *P*_true-lumen_ = 130 mmHg, the pressure in the experiments of Roach and colleagues on healthy porcine aortas. Change of *P*_tear-fl_ with changing axial stretch of the vessel is plotted in the inset. (b and c) Dependence of tear resistance, 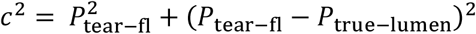, on changing tissue properties; *G*_*c*_ and *K*_t_/*K*_t0_ denote the characteristic energy of tearing and fold-change of tangent stiffness of the tissue. The solid lines are power-law fits to results of the numerical simulations. Changes of *P*_tear-fl_ with *G*_*c*_ and *K*_t_/*K*_*t*0_ are plotted in the insets. In panels (b) and (c), all axes of the main plots are logarithmic.(d) decreases with increases in the initial torn areaThe circles correspond to models with isotropic changes of the area, triangles correspond to elongated torn areas, and diamonds correspond to the area recorded during tear propagation in Fig. 2. Variation of with is shown in the inset. All axes in panel (d) were plotted on a logarithmic scale. See Fig S6 for additional results of a parametric study that confirms the power-law relations.

**Figure 5.**
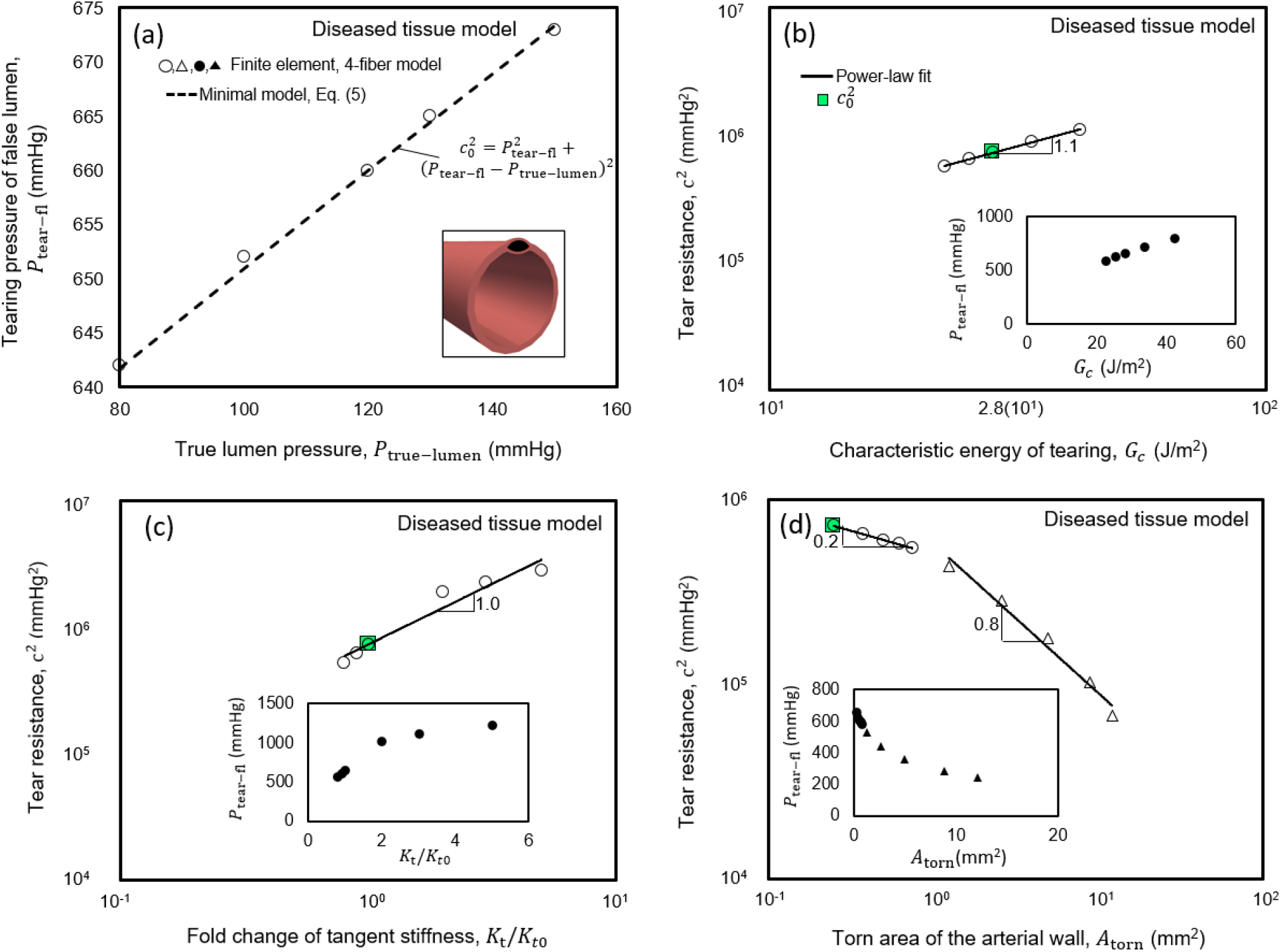
Dependencies of *P*_tear−fl_ on luminal pressure with tear properties assessed by the full nonlinear finite element model employing a Fung-type constitutive relation for a diseased human aorta. The numerical values and scaling relations are comparable to those for the healthy human aorta, presented in Fig. 4. The axes of main plots in panels (b-d) scale logarithmically.

### The pressure of tear progression reduces with increasing size of the initial torn area

To assess the effect of the initial area of tissue damage (due to the insertion of a needle or possible *in vivo* defects) on the pressure required for tear propagation, we performed similar finite element computations for the healthy human aorta using various values of *A*_torn_. In the first set of simulations, the torn area was prescribed isotropically (circles in Fig. 4(d)), whereas, in the second set, it had an elongated shape (triangles in Fig. 4(d)). In both cases, *P*_tear-fl_ decreased with increasing values of *A*_torn_. We also calculated the torn area as the tear propagated, starting from a small *A*_torn_, using the pressure values in Fig. 2. The exponent y (Eq. 5), which relates tear resistance to *A*_torn_, was evaluated as 0.5, 0.6, and 0.7 for cases with isotropic and elongated changes of *A*_torn_ and the case of propagation, starting from a small *A*_torn_. Again, similar trends were observed in computations for the diseased human aorta (Fig. 5). The dependency of *P*_tear-fl_ in response to combinations of changes of the tested parameters is further described in Supplementary Information (Fig. S6). Fig. S7 also shows possible effects on the pressure of tearing due to different values of the deposition stretch for collagen, though all other results herein assume a value of 1.08 for this important parameter consistent with widely assumed values. Again, the simulations were consistent with our validation experiments, which showed a decrease of *P*_tear-fl_ with increasing needle size (*p-value*=0.01) as the gauge changed from 30 to 25, thus increasing the size of the initial tear made by inserting the needle. The experimentally measured changes in *P*_tear-fl_ with *A*_torn_ also agreed qualitatively with the finite element model results. *P*_tear-fl_ changed 1.9-fold in comparing data for the 30- and 25-gauge needles.

### Tear propagation follows oblique regions of weakness of the wall

We next used the finite element model to assess possible deflections of the path of tear propagation by mural inhomogeneities. In the two-dimensional model with homogeneous wall properties, the tear consistently propagated circumferentially (Fig. 6(a)), which was independent of the structure of the finite element mesh (Fig. S8 in Supplemental information). It is noted further that the validation experiments on healthy porcine aortas similarly showed circumferential tears remaining within the same medial layers, not redirecting toward the lumen or adventitia (Figure S5). We thus simulated possible localized changes in wall composition / organization to model areas of focal aberrant material accumulation (strengthening) or degradation (weakening), and thus strong inhomogeneities, consistent with histologically observed local accumulations of mucoid material, calcification, or fibrotic tissue. Regions of weakening had values of tangent stiffness and characteristic energies of tearing that were reduced by a factor of three in comparison with the surrounding media. Regions of strengthening had tangent stiffnesses and characteristic energies of tearing five times that of the surrounding media. Simulations with either radially oriented weakening or strengthening inhomogeneities suggested that tear propagation remained unchanged in the presence of radially oriented inhomogeneities. The tears passed through these inhomogeneities and continued to propagate in the circumferential direction (Fig. 6(b) and (c)). Tests with oblique weakenings showed, however, that the path of propagation of a tear may be deflected by these inhomogeneities. Focal regions of weakness oriented at angles of roughly 15 degrees with respect to the circumferential direction were able to guide the tear either inward toward the true lumen or outward toward the outer surface of the vessel (Fig. 6(d) and (e)). A strong oblique inhomogeneity could similarly deflect the path of tear propagation (Fig. 6(f)). Taken together, these results demonstrate the possibility that the path of propagation of a tear can be influenced dramatically by inhomogeneities in wall properties.

**Figure 6.**
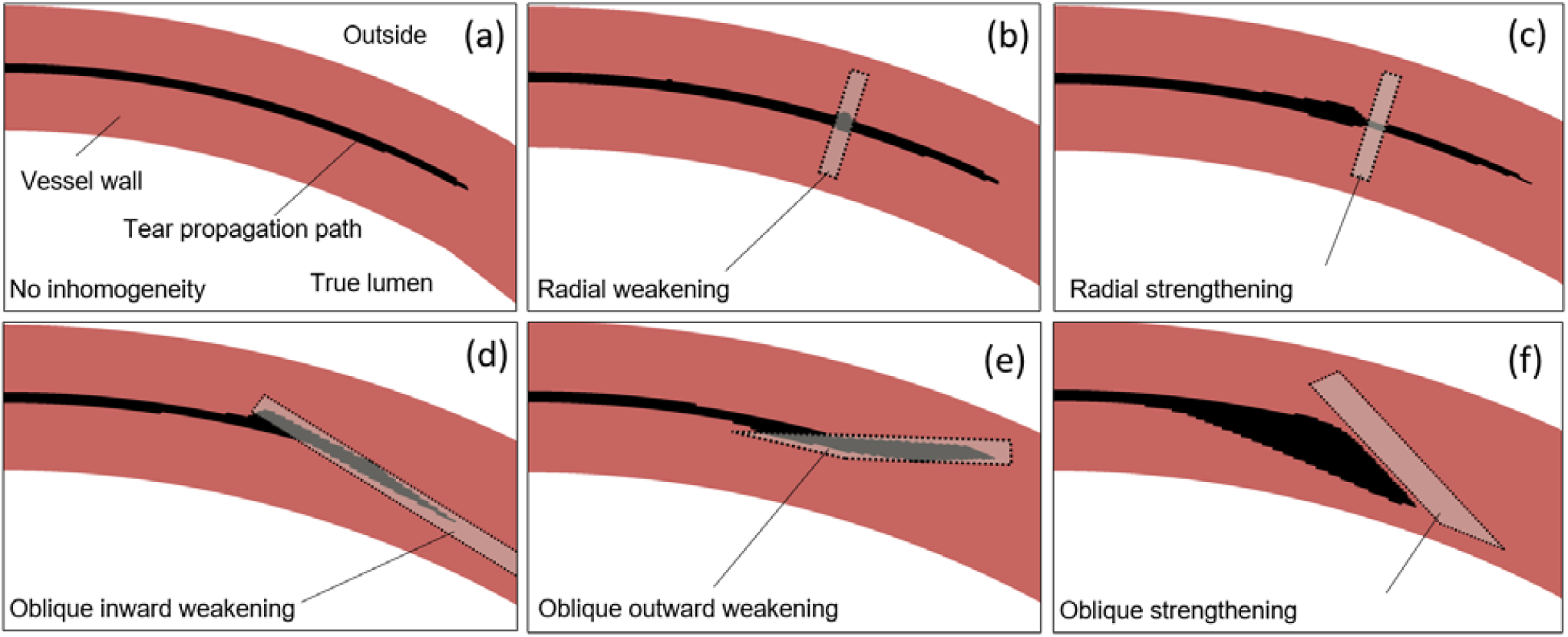
Inhomogeneous wall properties can direct the path of propagation of a tear away from the circumferential direction, guiding the tear toward either the true lumen or the outer surface of the vessel. Results from 2D finite element simulations shown (via post-processing) with color coding to highlight the undamaged tissue (dark), the prescribed inhomogeneity (light), and the tear (black). (a) The simulated tear tended to propagate circumferentially in the homogeneous anisotropic models of a healthy human aortic wall. (b and c) The tear similarly propagated circumferentially in the presence of either a radially oriented weakening or strengthening inhomogeneity. (d and e) An oblique region of weakness of the wall oriented at roughly 15-degree angles with respect to the circumferential direction could deflect the path of propagation toward either the true lumen or outside of the vessel. (f) A strengthening inhomogeneity oriented at 30 degrees with respect to the circumferential direction could also deflect the propagation of the tear. In panels (b), (c), and (f), there are build-ups of the injected fluid in the vicinity of the tear before the tear goes into a stiffer and stronger portion of the vessel wall. See Fig. S9 in Supplemental Information for similar findings for a model of a diseased human aortic wall.

## Discussion

Margot Roach and colleagues reported two particularly important experimental studies of aortic dissection in the 1990s but noted that the associated “mechanical analysis is complex.” ^13,30^ In both cases, they quantified the pressure required to initiate and propagate an intra-lamellar delamination within the porcine aorta independent of prior disease. We previously showed that a phase-field finite element model was able to capture the experimental findings reported in 1990 for excised traction-free slabs of aorta into which ink was injected under pressure.^4^ That study revealed differential mechanisms of intramural delamination in the thoracic and abdominal aorta as well as potential power-law relations that govern the propagation of a dissection. It was unclear, however, whether these relations depended strongly on the nearly flat *in vitro* geometry that was used experimentally and simulated computationally.

In this paper, we show that our basic phase-field finite element approach also captured similar data, reported by Roach and colleagues in 1999, for a more physiological geometry (cylindrical) and quasi-static biaxial loading (including the initial residual stress field). As in the Roach experiments on healthy porcine aortas, the simulated pressure required to initiate the tear was found herein for healthy and diseased human aortas to be much higher than that required to propagate the tear. The simulations revealed further that this pressure of tearing increases with acute increases in luminal pressure and axial stretch (Fig. 4(a), *inset*). Importantly, the *in vivo* value of axial stretch often diminishes in vessels having a propensity to dissect.^7^

Again, power-law relations emerged for the pressure at which the tear begins to propagate in terms of (proportionally) local wall stiffness and a characteristic energy of tearing and in terms of (inversely) the initial area of the tear. We submit that this power-law relation (Eq. 5), supported by simple linear models and confirmed via nonlinear finite element methods, helps to clarify and summarize what are otherwise complex coupled nonlinear effects and effectors. Such relations have promise to consolidate data from many different experiments on vessels having different vulnerabilities, a continuing experimental need. Finally, we also found that localized regions of intramural weakening or strengthening can affect the direction of the propagation, possibly turning an otherwise preferential circumferential propagation within a single lamellar unit (Fig. 6) towards either the lumen (possibly initiating a dissection or driving a re-entry tear) or the adventitia (possibly predisposing to transmural rupture). These findings point to the importance of improving *in vivo* imaging capability to assess risk of propagation of a pre-existing mural defect or dissection, particularly with regard to rupture potential. Collectively, the present results also confirm the importance of material properties as well as the type and size of the initial insult and the external loading conditions. Identifying precise mechanisms of pathogenesis yet remains challenging, though intimal tears^26,37^ and focal intramural accumulations of highly negatively charged mucoid material^23,32^ are potential initiators. At the nucleation site of the tear, when the size is below a millimeter, a normal static blood pressure is too low to separate the aortic layers and advance the tear. Although there is a need to study effects of hemodynamic loading, the Gibbs-Donnan swelling pressure due to mucoid materials may be sufficient to initiate the separation of the elastic lamella in such cases of micro-defects.^1,32^ With increasing volume of an intramural defect, the concentration of the charges may decrease, resulting in a reduction in swelling pressure. The inverse relation between the critical pressure required for tearing and the size of the false lumen suggests the possibility of the continuation of delamination of larger defects either by elevated blood pressure or the progressive accumulation of mucoid material at higher concentrations. Continuing focus on intra-lamellar structure and strength similarly promises to provide continued insight.^26,39^

Advantages of the present study include the qualitatively similar results obtained from nonlinear phase-field finite element models, linear analytical models and, consistent with the experiments of Roach and colleagues, simple confirmatory experiments on excised healthy porcine aortas. We focused our simulations, however, on models of the human aorta, both healthy and diseased, with geometries and pseudoelastic properties inferred from well-regarded experimental studies. Notwithstanding these advantages, there are limitations. We focused on potential dissections and thus initial defects within the media, without consideration of differential properties of the media and adventitia.^24^ The latter would be particularly important in simulations of transmural rupture, which were not considered herein. We also focused on energy-based calculations, not delineating possible effects of normal versus shear stresses in the damage process.^8^ In this regard, there is a need for additional experiments to contrast potential effects of normal and shear stresses in curved versus straight segments of the aorta; we considered only straight segments similar to the studies of Roach. Uncontrolled hypertension, vascular aging, and genetic mutations (e.g., leading to Marfan syndrome or Loeys-Dietz syndrome) often predispose to aortic dissections, and mouse models have revealed that each of these conditions associates with significant changes in aortic composition and material properties.^7,10,16^ There is also a need to repeat the experiments of Roach and colleagues, and then associated computational simulations, for vessels excised from vulnerable humans or animals that model these key risk factors. Although many to most aortic dissections occur independent of aneurysm,^21,27^ there is similarly a need to repeat the present analyses for different degrees of aneurysmal dilatation since they also associate with marked changes in aortic composition and properties, which can vary locally.^9^ We did not consider effects of aortic diameter herein. Perhaps the greatest need is to delineate experimentally the precise focal versus global changes in the wall that likely initiate dissection.^3^ Exquisite imaging of the microstructure of the vulnerable aorta followed by pressurization tests under *in vivo* conditions is likely the best way forward.

In summary, previous computational models (including those based on partition of unity and extended finite elements) have provided considerable insight into the biomechanics of aortic dissection^1,20,36,38,42^ as well as the importance of the effects of solid-fluid interactions.^5,14,25^ In this paper, we showed further that phase-field based finite element models can capture key experimental findings and reveal important new power-law relations that promise to aid in understanding further the biomechanical mechanisms of initiation and propagation of aortic dissection.

## Acknowledgments

We thank the Sinusas Lab at Yale University for providing us with aortic tissue samples. This work was supported, in part, by a grant from the US National Institutes of Health (UOl HLl425l8).

## Appendix: Pressure of tearing in analytical models of two adhered linearly elastic plates or membranes

In our previous study of region-specificity of dissections,^4^ we used mathematical relationships amongst critical pressure, stiffness, and characteristic energy of tearing proposed by Gent and Lewandowski^18^ to model delamination by injection of cut-open arterial segments. To obtain similar mathematical relations in the case of an inflated artery we built simplified models that extend the analysis of Gent and Lewandowski to include two flaps of tissue interleaved by two pools of pressurized fluid (Fig. Al and Fig. 3). The pressure of the false lumen deforms the two flaps of tissue while the pressure of the true lumen resists the deformation of the inner flap, with the mechanics of the four elements coupled.

Consistent with the computational models, we employed an energy-based approach^19^ to find the critical pressure required to separate two adhered linear elastic circular plates or membranes in the presence of a counter-acting fluid pressure acting over the outer surface of one of the circular elements (Fig. 3(b) and (c)). In these minimalistic models, we neglect the deformation of the artery outside of the circular areas. The solid bodies share the same radius *a*, thickness, *h*, elastic modulus, *E*, and Poisson’s ratio, *v*. The energy required to advance the torn surface by unit area is *G*_*c*_. The energy-based approach requires evaluation of the total energy that consists of the energies of the elastic deformation of the inner and outer flaps of tissue, *U*_*inner*_ and *U*_*outer*_, the energy loss by the pressure in the true and false lumens, −*P*_tl_ Δ*V*_tl_ and −*P*_fl_ Δ*V*_fl_, and finally the energy of tearing a *πa*^2^ *G*_*c*_. The volumes of the true and false lumens and the deformation of the flaps interact such that Δ*V*_fl_ = *V*_out-flap_ +*V*_in-flap_, with Δ*V* _tl_ = −*V*_in-flap_ · *V*_out-flap_ and *V* _in-flap_ are the volumes of fluid enclosed by the deformed outer and inner flaps. In comparison with the energy in the finite element implementation, instead of prescribing the injection volume, here we consider the state of deformation and tearing due to application of a constant pressure at the false lumen. The stationary point of the total energy with respect to the tear size determines the pressure of tearing.

**Figure A1.**
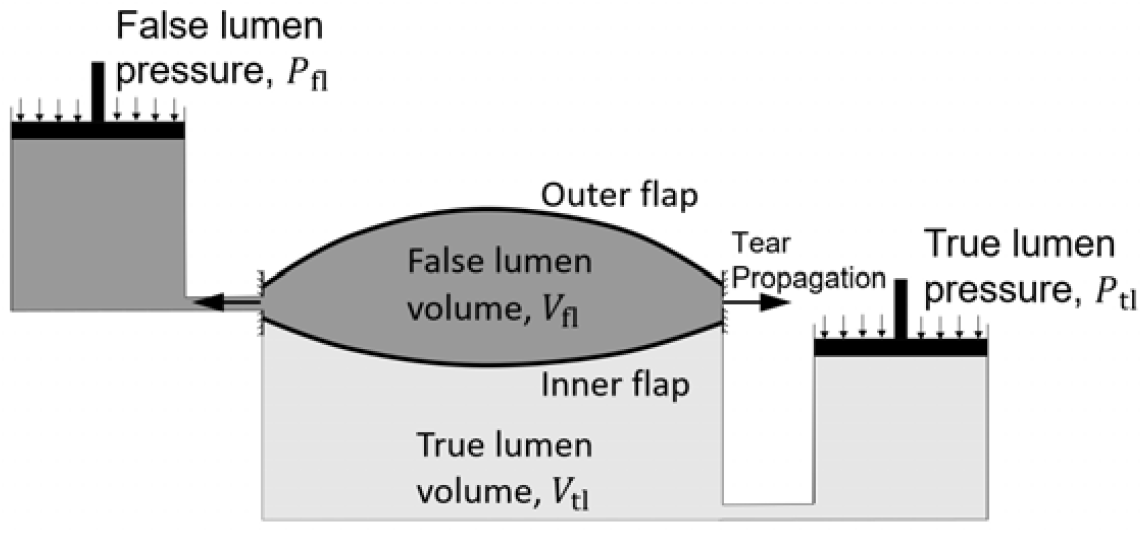
Schematic of the deformation of the inner and outer flaps of the aortic wall by the pressure of the true lumen and the intramural pressure of injection in the false lumen. The pressure of the false lumen deforms the outer flap of tissue while the differential of pressure of the true and false lumens deforms the inner flap. The interaction of the two pools of fluid is modeled by the loss of the volume of false lumen as the volume of the true lumen increases and vice versa.

In the case of the small deformation of a single linearly elastic plate, the axisymmetric displacement of a thin plate,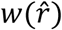, at a radius is governed by 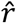. is the total uniformly distributed pressure and *B*=*Eh*^3^/(12(1−*v*^2^)) is the bending stiffness of the plate. Given the prescribed zero displacement at the periphery of the plate (Fig. 3(c)) and the non-zero displacement in the middle of the plate, 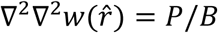. If we consider that the torn area is initially negligible, the energy required to produce a torn area of radius *a* is *E*_tear_=*πa*^2^ *G*_*c*_. The volume of fluid enclosed by the deforming plate is, 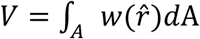 where *A* is the circular area of the plate before deformation. The elastically stored energy in the plate is *U*=*a*^6^*P*^2^π(1−*v*^2^)/(32*Eh*^3^).

The outside plate, located between the false lumen and the outside surface of the vessel, is loaded by the pressure from the false lumen, *P*=*P*_tear−fl_, whereas, the inside plate, sandwiched between the true and false lumens, is loaded by the differential of pressure of the two *P*=*P*_tear−fl_−*P*_true−lumen_. Following the approach of Griffith,^19^ the critical pressure of tearing corresponds to the tear size at the stationary point of *E*_total_, that is, where ∂*E*_total_/∂*a*=0. In the case of two-plates,

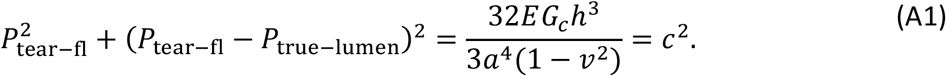

For the tear to advance between two deforming membranes (without bending stiffness) interleaved by two pools of fluid, we extend the analysis of Gent and Lewandowski^18^ for a single membrane. The change of potential of the fluid pressure is −*PV*. The deformation of the membrane may be characterized by approximate relations for the maximum deflection of the membrane, *δ*, and the volume of the fluid enclosed by the membrane: and *V* =*C*_1_ π *a*^2^δ. Since *P* ∝ *V*^3^, and *U*=∫_*v*_ *PdV, U*=*PV*/4 in case of the nonlinear deformation of the membrane. By repeating the same steps as in the case of two plates, we obtain the relationship

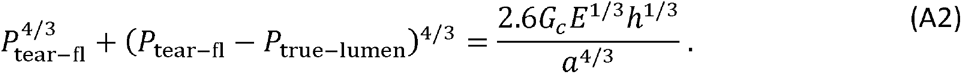

Importantly, both Eqs. (A1 and A2) suggest power-law relations for tear resistance, which motivated a new interpretation of nonlinear finite element results as shown in Figs. 4 and 5. Models with tense membranes and beams are presented in more detail in the Supplementary Information (Eqs. S4-S14), which similarly support this interpretation.

## References

1. Ahmadzadeh, H., M. K. Rausch, and J. D. Humphrey. Modeling lamellar disruption within the aortic wall using a particle-based approach. Scientific Reports 9:15320, 2019.

2. Alnæs, M., J. Blechta, J. Hake, A. Johansson, B. Kehlet, A. Logg, C. Richardson, J. Ring, M. E. Rognes, and G. N. Wells. The FEniCS Project Version 1.5. Archive of Numerical Software 3:, 2015.

3. Aslanidou, L., M. Ferraro, G. Lovric, M. R. Bersi, J. D. Humphrey, P. Segers, B. Trachet, and N. Stergiopulos. Co-localization of microstructural damage and excessive mechanical strain at aortic branches in angiotensin-II-infused mice. Biomech Model Mechanobiol 19:81–97, 2020.

4. Ban, E., C. Cavinato, and J. D. Humphrey. Differential propensity of dissection along the aorta. Biomech Model Mechanobiol 20:895–907, 2021.

5. Bäumler, K., V. Vedula, A. M. Sailer, J. Seo, P. Chiu, G. Mistelbauer, F. P. Chan, M. P. Fischbein, A. L. Marsden, and D. Fleischmann. Fluid–structure interaction simulations of patient-specific aortic dissection. Biomech Model Mechanobiol 19:1607–1628, 2020.

6. de Beaufort, H. W. L., A. Ferrara, M. Conti, F. L. Moll, J. A. van Herwaarden, C. A. Figueroa, J. Bismuth, F. Auricchio, and S. Trimarchi. Comparative Analysis of Porcine and Human Thoracic Aortic Stiffness. European Journal of Vascular and Endovascular Surgery 55:560–566, 2018.

7. Bellini, C., M. R. Bersi, A. W. Caulk, J. Ferruzzi, D. M. Milewicz, F. Ramirez, D. B. Rifkin, G. Tellides, H. Yanagisawa, and J. D. Humphrey. Comparison of 10 murine models reveals a distinct biomechanical phenotype in thoracic aortic aneurysms. Journal of The Royal Society Interface 14:20161036, 2017.

8. Bellini, C., N. J. Kristofik, M. R. Bersi, T. R. Kyriakides, and J. D. Humphrey. A hidden structural vulnerability in the thrombospondin-2 deficient aorta increases the propensity to intramural delamination. Journal of the Mechanical Behavior of Biomedical Materials 71:397–406, 2017.

9. Bersi, M. R., C. Bellini, J. D. Humphrey, and S. Avril. Local variations in material and structural properties characterize murine thoracic aortic aneurysm mechanics. Biomech Model Mechanobiol 18:203–218, 2019.

10. Bersi, M. R., R. Khosravi, A. J. Wujciak, D. G. Harrison, and J. D. Humphrey. Differential cell-matrix mechanoadaptations and inflammation drive regional propensities to aortic fibrosis, aneurysm or dissection in hypertension. Journal of The Royal Society Interface 14:20170327, 2017.

11. Bourdin, B., G. A. Francfort, and J.-J. Marigo. The Variational Approach to Fracture. J Elasticity 91:5–148, 2008.

12. Campbell, J. D. On the theory of initially tensioned circular membranes subjected to uniform pressure. The Quarterly Journal of Mechanics and Applied Mathematics 9:84–93, 1956.

13. Carson, M. W., and M. R. Roach. The strength of the aortic media and its role in the propagation of aortic dissection. Journal of Biomechanics 23:579–588, 1990.

14. Dillon-Murphy, D., A. Noorani, D. Nordsletten, and C. A. Figueroa. Multi-modality image-based computational analysis of haemodynamics in aortic dissection. Biomech Model Mechanobiol 15:857–876, 2016.

15. Dingemans, K. P., P. Teeling, A. C. van der Wal, and A. E. Becker. Ultrastructural pathology of aortic dissections in patients with Marfan syndrome: Comparison with dissections in patients without Marfan syndrome. Cardiovascular Pathology 15:203–212, 2006.

16. Ferruzzi, J., D. Madziva, A. W. Caulk, G. Tellides, and J. D. Humphrey. Compromised mechanical homeostasis in arterial aging and associated cardiovascular consequences. Biomech Model Mechanobiol 17:1281–1295, 2018.

17. García-Herrera, C. M., D. J. Celentano, M. A. Cruchaga, F. J. Rojo, J. M. Atienza, G. V. Guinea, and J. M. Goicolea. Mechanical characterisation of the human thoracic descending aorta: experiments and modelling. Computer Methods in Biomechanics and Biomedical Engineering 15:185–193, 2012.

18. Gent, A. N., and L. H. Lewandowski. Blow-off pressures for adhering layers. Journal of Applied Polymer Science 33:1567–1577, 1987.

19. Griffith, A. A. VI. The phenomena of rupture and flow in solids. Philosophical Transactions of the Royal Society of London. Series A, Containing Papers of a Mathematical or Physical Character 221:163–198, 1921.

20. Gültekin, O., S. P. Hager, H. Dal, and G. A. Holzapfel. Computational modeling of progressive damage and rupture in fibrous biological tissues: application to aortic dissection. Biomech Model Mechanobiol 18:1607–1628, 2019.

21. Guo, D.-C., E. S. Regalado, C. Minn, V. Tran-Fadulu, J. Coney, J. Cao, M. Wang, R. K. Yu, A. L. Estrera, H. J. Safi, S. S. Shete, and D. M. Milewicz. Familial Thoracic Aortic Aneurysms and Dissections. Circulation: Cardiovascular Genetics 4:36–42, 2011.

22. Hughes, T. J. R. The Finite Element Method: Linear Static and Dynamic Finite Element Analysis. Courier Corporation, 2012, 706 pp.

23. Humphrey, J. D. Possible Mechanical Roles of Glycosaminoglycans in Thoracic Aortic Dissection and Associations with Dysregulated TGF-β. J Vasc Res 50:1–10, 2013.

24. Kawamura, Y., S.-I. Murtada, F. Gao, X. Liu, G. Tellides, and J. D. Humphrey. Adventitial remodeling protects against aortic rupture following late smooth muscle-specific disruption of TGFβ signaling. Journal of the Mechanical Behavior of Biomedical Materials 116:104264, 2021.

25. Keramati, H., E. Birgersson, J. P. Ho, S. Kim, K. J. Chua, and H. L. Leo. The effect of the entry and re-entry size in the aortic dissection: a two-way fluid–structure interaction simulation. Biomech Model Mechanobiol 19:2643–2656, 2020.

26. Pal, S., A. Tsamis, S. Pasta, A. D’Amore, T. G. Gleason, D. A. Vorp, and S. Maiti. A mechanistic model on the role of “radially-running” collagen fibers on dissection properties of human ascending thoracic aorta. Journal of Biomechanics 47:981–988, 2014.

27. Pape, L. A., T. T. Tsai, E. M. Isselbacher, J. K. Oh, P. T. O’Gara, A. Evangelista, R. Fattori, G. Meinhardt, S. Trimarchi, E. Bossone, T. Suzuki, J. V. Cooper, J. B. Froehlich, C. A. Nienaber, and K. A. Eagle. Aortic Diameter ≥5.5 cm Is Not a Good Predictor of Type A Aortic Dissection. Circulation 116:1120–1127, 2007.

28. Pasta, S., J. A. Phillippi, T. G. Gleason, and D. A. Vorp. Effect of aneurysm on the mechanical dissection properties of the human ascending thoracic aorta. The Journal of Thoracic and Cardiovascular Surgery 143:460–467, 2012.

29. Rivlin, R. S., and A. G. Thomas. Rupture of rubber. I. Characteristic energy for tearing. Journal of Polymer Science 10:291–318, 1953.

30. Roach, M. R., J. C. He, and R. G. Kratky. Tear propagation in isolated, pressurized porcine thoracic aortas. Can J Cardiol 15:569–575, 1999.

31. Roach, M. R., and S. H. Song. Variations in strength of the porcine aorta as a function of location. Clin Invest Med 17:308–318, 1994.

32. Roccabianca, S., G. A. Ateshian, and J. D. Humphrey. Biomechanical roles of medial pooling of glycosaminoglycans in thoracic aortic dissection. Biomech Model Mechanobiol 13:13–25, 2014.

33. Roccabianca, S., C. A. Figueroa, G. Tellides, and J. D. Humphrey. Quantification of regional differences in aortic stiffness in the aging human. Journal of the Mechanical Behavior of Biomedical Materials 29:618–634, 2014.

34. Sherifova, S., and G. A. Holzapfel. Biomechanics of aortic wall failure with a focus on dissection and aneurysm: A review. Acta Biomaterialia 99:1–17, 2019.

35. Sommer, G., S. Sherifova, P. J. Oberwalder, O. E. Dapunt, P. A. Ursomanno, A. DeAnda, B. E. Griffith, and G. A. Holzapfel. Mechanical strength of aneurysmatic and dissected human thoracic aortas at different shear loading modes. Journal of Biomechanics 49:2374–2382, 2016.

36. Thunes, J. R., J. A. Phillippi, T. G. Gleason, D. A. Vorp, and S. Maiti. Structural modeling reveals microstructure-strength relationship for human ascending thoracic aorta. Journal of Biomechanics 71:84–93, 2018.

37. Tong, J., Y. Cheng, and G. A. Holzapfel. Mechanical assessment of arterial dissection in health and disease: Advancements and challenges. Journal of Biomechanics 49:2366–2373, 2016.

38. Wang, L., S. M. Roper, N. A. Hill, and X. Luo. Propagation of dissection in a residually-stressed artery model. Biomech Model Mechanobiol 16:139–149, 2017.

39. Wang, R., X. Yu, and Y. Zhang. Mechanical and structural contributions of elastin and collagen fibers to interlamellar bonding in the arterial wall. Biomech Model Mechanobiol 20:93–106, 2021.

40. Weinsaft, J. W., et al. Aortic Dissection in Patients With Genetically Mediated Aneurysms: Incidence and Predictors in the GenTAC Registry. Journal of the American College of Cardiology 67:2744–2754, 2016.

41. Williams, M. L. The continuum interpretation for fracture and adhesion. Journal of Applied Polymer Science 13:29–40, 1969.

42. Yu, X., B. Suki, and Y. Zhang. Avalanches and power law behavior in aortic dissection propagation. Science Advances 6:eaaz1173, 2020.

