## Supplementary material for "Critical Pressure of Intramural Delamination in Aortic Dissection": SI

#### Supplementary Methods

##### Prescription of the initial torn area

A damaged region was initialized in the model by prescribing the phase-field damage variable $\phi = 1$ therein. This damaged region represented the initial incision due to insertion of the needle. This region created an initial torn surface, $A_{\mathrm{torn}}$, of area $l_z \times l_\theta$ within the wall (Fig. S1). $l_z$ and $l_\theta$ represent the length of the torn area in the axial and circumferential directions, respectively. The values of $l_z$ and $l_\theta$ were changed in simulations of varying torn areas.

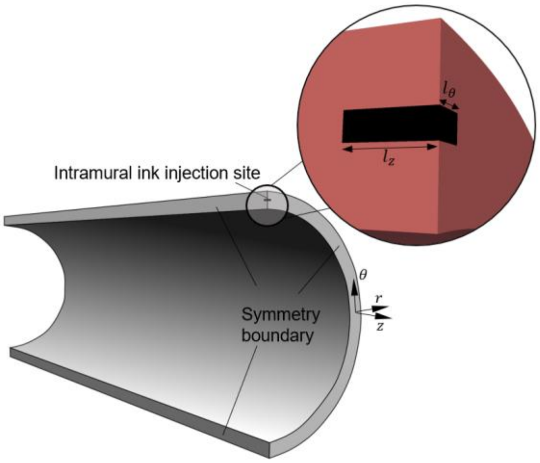


**Figure S1. Parameters controlling the size of the injection pool, that is, the initial damage volume.** It is useful to focus on the torn area since the energy-based solution accounts for the energy required to create new surfaces.

#### Additional details on the variational phase-field finite element model

Our implementation of a phase-field finite element model having a fully nonlinear arterial constitutive relation is outlined in the main text and described in large part in our previous publication.[1] The present section complements the description in the main text. Referring to Eq. (1) therein, the energy of deformation $E_{\text{deformation}}$ is

$$E_{\text{deformation}} = \int_V \left( ((1-\phi)^2 + \varepsilon)\, \alpha_{K_t} W_{\text{wall}} + W_{\text{vol}} \right) dV, \qquad \text{(S1)}$$

where $\phi \in [0,1]$ is the damage variable and $\varepsilon$ is a small number that sets a minimum energy and stiffness for the fully damaged tissue that facilitates numerical calculations. The $\alpha_{K_t}$ is a pre-factor that we used to vary tissue tangent stiffness in studies of the effect of tissue tangent stiffness on aortic tearing. $W_{\text{wall}}$ is the strain energy density (per unit volume) of the arterial wall that depends on values of the principal stretches. Toward this end, the deformation gradient was multiplicatively decomposed into deformation and deposition parts. The deformation portion was determined as part of the solution of the finite element model, while the deposition portion was prescribed. Importantly, residual stresses can arise naturally when prescribing non-unity deposition stretches within the strain energy.[10] In the case of healthy tissue, these deposition stretches were $G^e = 1.2$ and $G^k = 1.08$ for elastin and collagen, respectively. $G^e$ actually describes both the true deposition stretch in development and the *in vivo* prestretch that arises during somatic growth since elastin does not turnover in maturity. $G^k$ represents only the deposition stretch for each of the four families of collagen fibers turnover in maturity. Our pseudoelastic constitutive model thus represents the deformation of tissue with respect to a reference (loaded) *in vivo* state.

In Eq. (S1), $W_{\text{vol}}$ is an expression used to enforce the incompressibility of tissue

$$W_{\text{vol}} = -(1-\phi)^3\, p\, (\lambda_1 \lambda_2 \lambda_3 - 1) - \frac{p^2}{2\epsilon}. \qquad \text{(S2)}$$

Minimization of the first term on the right-hand side with respect to $p$ ensures incompressibility. Incompressibility is not enforced for damaged tissue, and the pre-factor $(1-\phi)^3$ ensures a faster degradation of this condition compared with strain energy.[8] The second term on the right-hand side regularizes the solution of $p$ in the damaged tissue, in an approach similar to the perturbed Lagrangian method.[14] $\epsilon$ is a large number.

The energy of tearing was expressed using the Ambrosio-Tortorelli I[11] expression

$$E_{\text{tear}} = \frac{3\alpha_{G_c} G_c}{8} \int_V \left( \frac{\phi}{l} + l |\nabla \phi|^2 \right) dV. \qquad \text{(S3)}$$

$G_c$ is the energy expended to extend the torn area by one unit. Here, $\alpha_{G_c}$ is a pre-factor that was used to vary $G_c$ to assess the effect of changing $G_c$ on aortic tearing. $l$ controls the length-scale of the decay of the damage field, chosen on the same order as the finite element mesh cell size, consistent with previous three-dimensional models of fracture.[2]

Note that the progress of injection was controlled by incrementally prescribing the volume of injection. When $E_{\text{pressurized−fluid}}$, introduced in Eq. (2) of the main text, is minimized with respect to the global Lagrange multiplier, $m$, the total volume of the injected fluid, or damaged tissue in the deformed configuration, is prescribed as $V_{\text{injection}}$. Here, $\int_V \phi\lambda_1\lambda_2\lambda_3\, dV$ represents the total volume of damaged tissue in the deformed configuration. This method is our contribution to the computational approach and replaces the use of displacement or pressure boundary conditions for applying loads. It enables tear propagation to be modeled past the maximum pressure point while avoiding computational instabilities.

#### Finite element mesh convergence study

We analyzed the sensitivity of the critical pressure of tearing ($P_{\text{tear−fl}}$) to the size of the cells in the finite element mesh (Fig. S2). The number of elements per wall thickness was quantified by passing a radial line through the wall thickness and counting the number of elements crossed by the line.

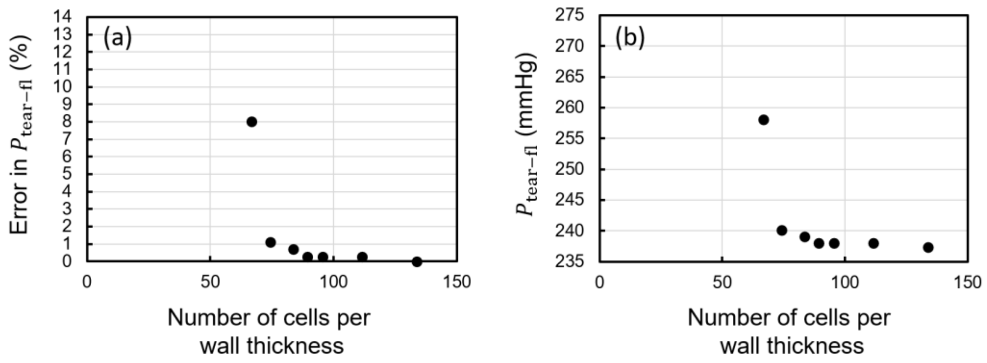


**Figure S2. Dependence of the pressure of tearing on the size of the cells in the finite element mesh.** The error is quantified relative to the pressure of tearing in the finest tested mesh. The number of cells per wall thickness is calculated as the number of cells that will be crossed by a radial line going through the mesh of the vessel wall.

#### Aortic constitutive parameters

The mechanical behavior of diseased human aortic tissue was modeled by fitting the parameters of the same four-fiber family constitutive model to previously reported uniaxial and simple shear stress-strain data from diseased (aneurysmal) aortas, previously applied to computationally model dissections.[7] Previous reports from our group indicated a decrease in the *in vivo* axial stretch in models of dissection. Hence, we reduced by a similar factor the axial and deposition stretches in the diseased material model.

In the published results on the energy required for tearing arterial tissue, we did not find a clear relation between diseases and energy of tearing. Hence, we did not change this parameter. We also used arterial radius and thickness reported previously.[7] The parameters for both healthy and diseased tissue are listed in Table S1.

**Table S1. Values of the material parameters within the four-fiber model that fit existing experimental results on healthy and diseased human aorta.**

| | $\bar{c}$ (kPa) | $c_1^1$ (kPa) | $c_2^1$ | $c_1^2$ (kPa) | $c_2^2$ | $c_1^{3,4}$ (kPa) | $c_2^{3,4}$ | $\alpha_0$ | $R_{0-\text{in}}$ (mm) | $h$ (mm) | Ref. |
|---|---|---|---|---|---|---|---|---|---|---|---|
| Healthy | 24.7 | $10^{-5}$ | $4(10^{-5})$ | $5(10^{-5})$ | $2(10^{-5})$ | 90.1 | 5.3 | 0.74 | 11.75 | 1.75 | 10 |
| Diseased | 53.2 | 20.8 | 0.5 | 53.2 | 18.7 | 9.8 | 0.5 | 1.57 | 15.0 | 2.5 | 7 |

#### Prescription of material inhomogeneities

Inhomogeneities of the vessel wall were prescribed by defining the aforementioned pre-factors $\alpha_{K_t}$ and $\alpha_{G_c}$ as spatially varying fields, included as mesh functions (Fig. S3). In the case of the homogeneous wall, $\alpha_{K_t} = 1$ and $\alpha_{G_c} = 1$ everywhere. In case of an inhomogeneity, $\alpha_{K_t}$ and $\alpha_{G_c}$ equaled 1 everywhere except for the area of the inhomogeneity. They were larger or smaller than 1 in cases of strengthenings and weakenings within the wall, respectively.

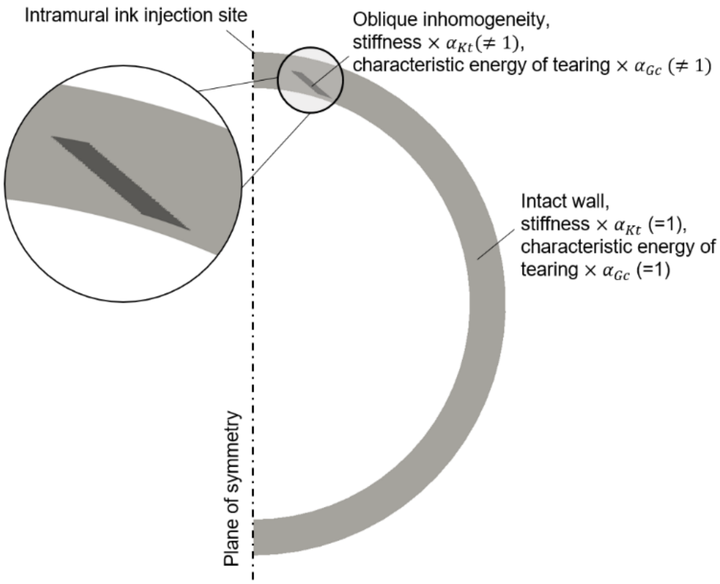


**Figure S3. The mesh functions used for mapping inhomogeneous material properties.** The spatial fields $\alpha_{Kt}$ and $\alpha_{Gc}$ were 1 in the intact region of the arterial wall; values larger and smaller than 1 were used to model local strengthenings and weakenings, of the wall, respectively.

#### Further details on the analytical models: cases with two beams and highly tensed membranes

If we consider two elastic beams with rectangular sections, instead of two plates, and repeat the approach of Appendix A of the main text, with vanishing deflection and angles at the end of the beams (Fig. S4), we arrive at a relationship similar to Eq. (A1) of the main text. We briefly describe the derivation of this relationship here.

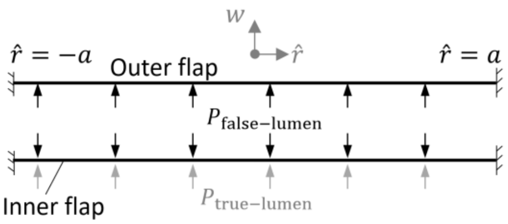


**Figure S4. Minimal analytical model with two beams and two pools of fluid.**

The bending of a beam with bending stiffness $EI$ deflected by a distributed load $P$ obeys[3]

$$\frac{\partial^4 w}{\partial \hat{r}^4} = \frac{P}{EI}, \tag{S4}$$

where $w$ is the lateral deflection of the beam and $\hat{r}$ is the coordinate along the length of the beam. By imposing the boundary conditions

$$w(a) = 0, w(-a) = 0, w'(a) = 0, w'(-a) = 0, \tag{S5}$$

where $w'$ denotes the derivative of $w$ with respect to $\hat{r}$, we find that

$$w(\hat{r}) = \frac{a^4 P}{24EI} - \frac{a^2 P \hat{r}^2}{12EI} + \frac{P \hat{r}^4}{24EI}. \tag{S6}$$

The volume enclosed by a bent beam is given by

$$V = 2b \int_0^a w(\hat{r})\, d\hat{r} = b\frac{2a^5 P}{45EI}, \tag{S7}$$

where $a$ is half of the length of the beam and $b$ is its depth out of the $w - \hat{r}$ plane. The elastic energy stored in a beam is

$$U = \frac{1}{2}(2)b \int_0^a P\, w(\hat{r})\, d\hat{r} = b\frac{a^5 P^2}{45EI} \tag{S8}$$

In the case with two beams, change of volume of the false lumen equals the sum of the volumes enclosed by the two flaps

$$\Delta V_{\mathrm{false-lumen}} = V_{\mathrm{outer-flap}} + V_{\mathrm{inner-flap}}, \tag{S9}$$

and change of volume of the true lumen is

$$\Delta V_{\text{true-lumen}} = -V_{\text{inner-flap}}. \tag{S10}$$

We note that the outer flap is loaded by $P_{\text{false-lumen}}$ while the inner flap is deformed by $P_{\text{false-lumen}} - P_{\text{true-lumen}}$. Total energy of the two flaps, the pistons providing pressure and the progression of tearing is

$$E_{\text{total}} = U_{\text{outer-flap}} + U_{\text{inner-flap}} - P_{\text{false-lumen}}\Delta V_{\text{false-lumen}} - P_{\text{true-lumen}}\Delta V_{\text{true-lumen}} + 2abG_c \tag{S11}$$

Following a Griffith-type approach[5,6,13], the stationary point of energy with respect to tear size determines the onset of the unstable propagation of the tear. Therefore, the solution of

$$\frac{\partial E_{\text{total}}}{\partial a} = 0 \tag{S12}$$

determines the pressure of tearing as

$$P_{\text{tear-fl}}^2 + (P_{\text{tear-fl}} - P_{\text{true-lumen}})^2 = \frac{3G_c E h^3}{2a^4} = c^2. \tag{S13}$$

Similarly, a linear solution for highly tensed membranes[4] results in the relationship

$$P_{\text{tear-fl}}^2 + (P_{\text{tear-fl}} - P_{\text{true-lumen}})^2 = \frac{8G_c h\sigma}{a^2} = c^2, \tag{S14}$$

where $\sigma$ is the initial stress in the membranes. Again, therefore, consistent with results in Appendix A of the main text, we see power-law relations emerge, which guided the interpretation of the finite element results.

#### A thought experiment to reduce our analytical models to previous models of single pools and single polymer films

Finally, consider a thought experiment that relates our models with two pools of fluid and two flaps of tissue (Fig. A1 of the main text) to the cases studied by Gent and Lewandowski[5] and Williams[13], which have a single polymer film and a single pool of fluid. First, we set $P_{\text{true-lumen}} = 0$. Next, we insert a stiff thin substrate between the inner and outer flaps. We adhere the flaps to the substrate at both ends. By this procedure, we reduce our system to two replicas of the system studied in previous theoretical models. Then the formulas from our model with two flaps reduce to $2P_{\text{tear-fl}}^{\eta} = G_c f(E, a)$. Here, $f$ is a function of $E$ and $a$, which is independent of $G_c$. We note that in the thought experiment, $G_c$ is the energy required to detach two flaps from the substrate. In comparison with the experiments with one flap and one pool of fluid, the $G_c$ here is twice the characteristic energy of tearing for the detachment of each film from the substrate, $G_c'$. In other words, $G_c' = 1/2G_c$. Therefore, the formulas reduce to $2P^{\eta}_{\text{tear-fl}} = 2G_c' f(E, a)$, and Eqs. (A1 and 2) of the main text and (S13) here compare directly with the previous relationships by Gent and Lewandowski[5] and Williams[13], thus providing another check.

#### Experimental validation studies

Figure S5 illustrates the experimental setup for the intramural injection tests on healthy excised porcine aortas. Perivascular tissue was removed by gentle dissection and all branches were ligated using nylon sutures. The samples were secured with ligatures on custom-drawn acrylonitrile butadiene styrene cannulas, mounted within a custom biaxial testing system, and immersed in Hanks' Balanced Salt Solution (HBSS) throughout the duration of the test. The samples were then extended axially to a stretch ratio of either 1.1 or 1.3, which was prescribed in our computational simulations. Following a short period of cyclic preconditioning (pressurization from 10 to 130 mmHg with the vessel held fixed at the axial stretch), the true lumen of the vessel was maintained at a pressure of 130 mmHg, equivalent to the value prescribed by Roach and colleagues.[9]

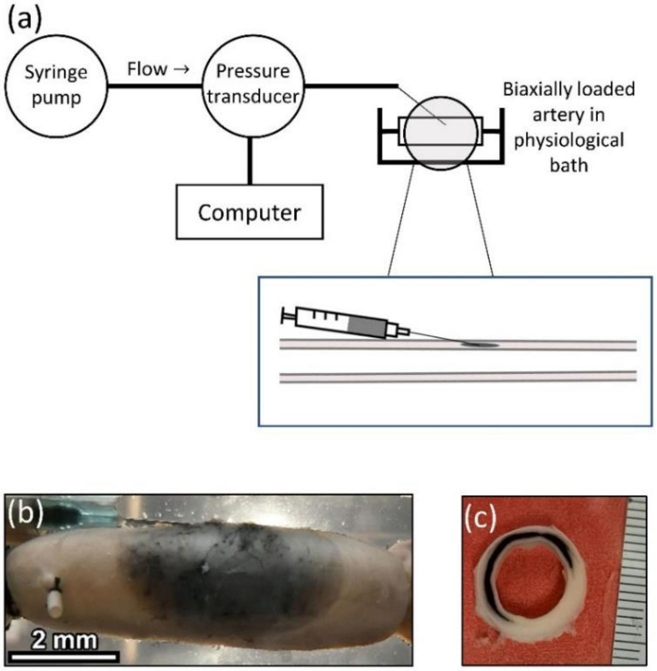


**Figure S5. Experimental setup for intramural injection of healthy cylindrical segments of porcine aorta.** (a) Schematic of the injection system and the sample. (b) Representative sample captured during the progression of the dissection induced by intramural injection, and (c) cross-section of the sample proximal to the needle insertion position cut after the test. Note the circumferential delamination.

Specifically, a 1-inch long 25G or 30G needle was inserted into the aortic wall in the direction of the longitudinal axis of the artery and at the minimum allowed angle to the external surface of the sample. The penetration depth was consistently 1 mm on the basis of the protruding length of the needle. In tests using the thicker needle, following insertion of the needle, we applied a small backward motion to withdraw the needle slightly to increase the initial volume of the damaged tissue while holding the end of the needle in the medial layer. A syringe pump (Thermo Fisher Scientific) was used to inject HBSS infused with India ink at the rate of 1 ml/min through a silicone pump tubing of 25 cm length and 2.79 mm inner diameter (Ismatec SA) connected at the opposite end to a 1-inch long 25- or 30-gauge needle (BD PrecisionGlide). A calibrated pressure transducer (Sensotec Inc.) was connected to the tubing in the proximity of the needle. The samples were randomly assigned to the 25G or 30G group. Following testing, a ring of each sample was sectioned to verify consistent dissection location. Statistical analysis of the data included comparison of the pressures of tearing for two comparable groups (same stretch, two different needles or same needle, two different stretches) using the Student's t-test.

### Supplementary Results

#### Variation of the pressure of tearing with simultaneous changes of parameters

To test the utility of the discovered power-law relations, we used the computational model to evaluate $P_{\text{tear-fl}}$ for different combinations of parameters. The parameter sets consisted of all possible combinations of the parameter values resulting from multiplying the baseline values of $G_c$, $K_t$, and $A_{\text{torn}}$ by elements of the sets {0.9, 1.0, 1.2}, {1, 1.2}, and {1, 2}, respectively. The resulting tear resistance, $c^2$, and pressure of tearing, $P_{\text{tear-fl}}$, obtained from the two approaches were comparable (Fig. S6).

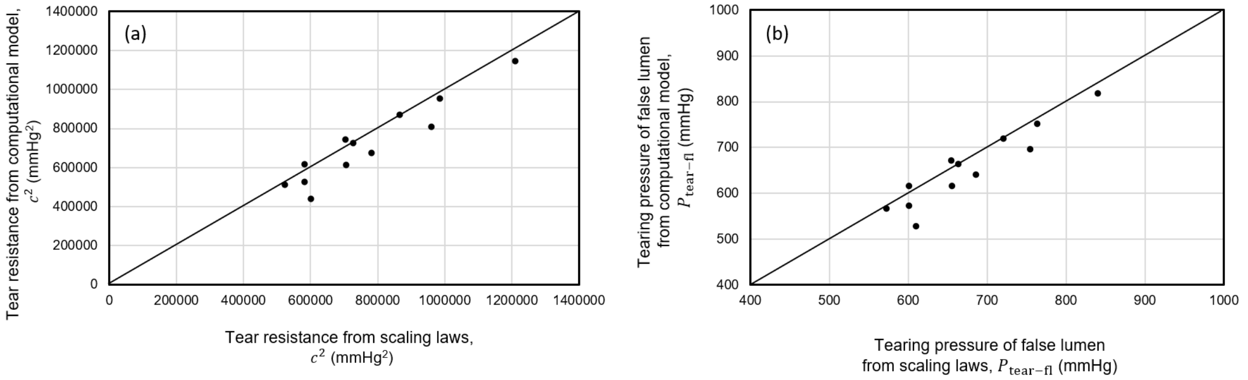


**Figure S6. Comparison of (a) tear-resistance and (b) intramural tearing pressure of false lumen between the computational model and the scaling laws with different combinations of the parameters.** The parameter sets consisted of all possible combinations of the parameter values resulting from multiplying the baseline values of $G_c$, $K_t$, and $A_{\text{torn}}$ by elements of the sets {0.9, 1.0, 1.2}, {1, 1.2}, and {1, 2}, respectively.

#### Influence of collagen deposition stretches on the tearing pressure

The four-fiber constitutive relation can be augmented with the introduction of deposition stretches, which account for the pre-stress that is built into different structurally significant constituents upon their incorporation within extant matrix. Associated constituent-specific pre-stresses, in turn, affect the residual stress field within the traction-free configuration. Here we show that the specific value of the collagen deposition stretch affects the phase-field based calculation of the pressure of tearing (Fig. S7), though we used a value of 1.08 in all other simulations, a value commonly used.

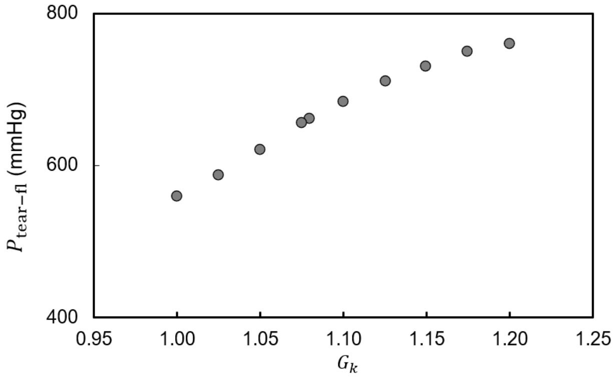


**Figure S7. Sensitivity of the pressure of tearing to the value of the deposition stretch for collagen.** Dependence of the finite element calculated value of $P_{\text{tear-fl}}$ as a function of the prescribed deposition stretch for collagen, $G_k$. A value of 1.08 was used in all other simulations herein.

#### Comparison of grid-based and non-grid-based meshes

We assessed the dependency of the direction of tear propagation on finite element mesh anisotropy. The original meshes were generated based on a grid and using an in-house code, generating non-uniformly refined meshes. We also generated non-grid-based triangular meshes using CGAL.[12] Using the same computational code as in the case of the grid-based mesh, we consistently observed circumferential propagation of tears (Fig. S8).

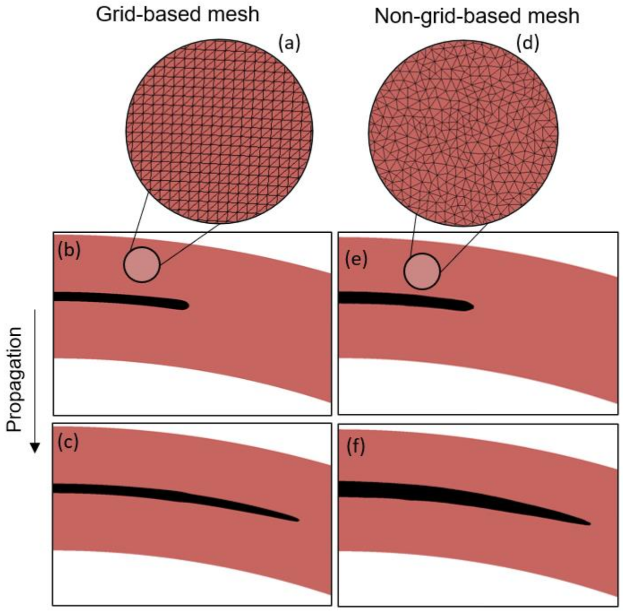


**Figure S8. Insensitivity of the circumferential propagation of injected fluid to mesh anisotropies.**

#### Roles of inhomogeneities in redirecting the path of tear propagation in diseased tissue

Effects of inhomogeneities on the propagation path of tears were also assessed using the diseased material model. Tears propagated circumferentially in the absence of inhomogeneities, similar to the case of healthy tissue. Oblique weakenings were able to divert the path of propagation into the true lumen (Fig. S9). These results suggest that local inhomogeneities can redirect tear propagation either toward the lumen or outside of the vessel. Here, in the absence of inhomogeneities, we observed circumferential propagation using multiple sets of material parameters for a particular constitutive relation. In the absence of exhaustive studies using different relations and parameters, the present computations do not rule out the possibility of tears propagating out of the vessel for other homogeneous material models that neglect the laminar microstructure.

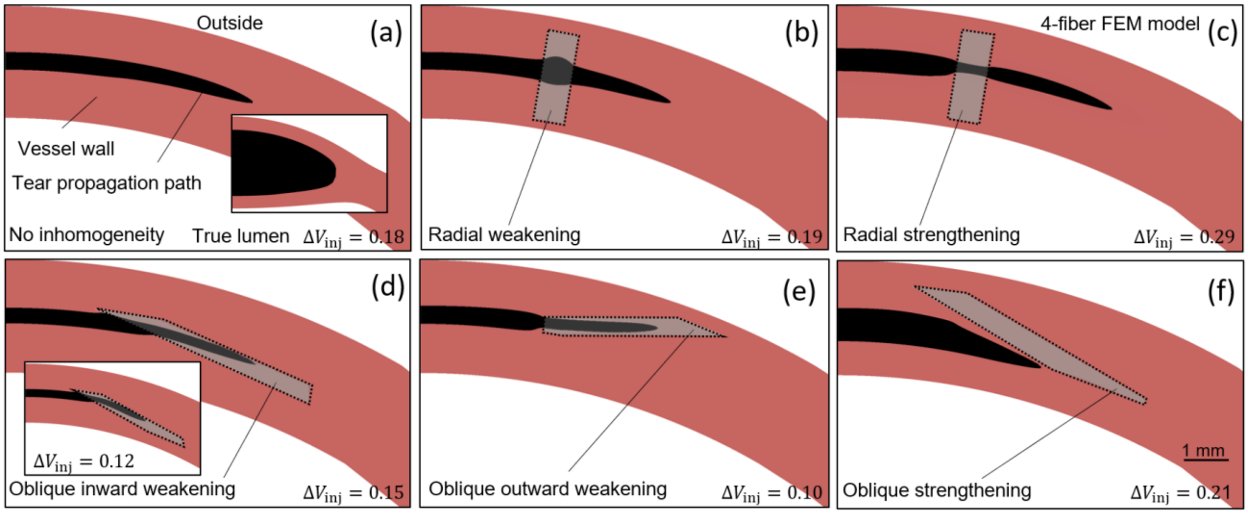


**Figure S9. Deviation of the path of tear propagation by inhomogeneities in the model of the diseased artery.** Results from 2D finite element computations using the four-fiber Fung-type constitutive model. The undamaged tissue is dark, the prescribed inhomogeneity is light, and the tear is black. Similar results for the model of a healthy artery were shown in Fig. 6 of the main text. The strengthening and weakenings were modeled by regions with $G_c$ and $K_t$ that were 3 or 1/3 times the rest of the wall, except for in the inset of panel (d), where a factor of 1/6 was used. $\Delta V_{\text{inj}}$ is the change in the amount of the injected fluid (in $mm^2$) normalized by the area of the undeformed aortic wall. The meshes and mesh functions in this model were similar to those used in the healthy model, illustrated in Figs. S8(a) and S3. All snapshots were taken in the unloaded configurations. The inset of panel (a) shows an example of the damaged wall in the loaded configuration (a). The deformed snapshot is comparable with the 3D models in Fig. 2 of the main text.

#### References


1. Ban, E., C. Cavinato, and J. D. Humphrey. Differential propensity of dissection along the aorta. *Biomech Model Mechanobiol* 20:895–907, 2021.

2. Borden, M. J., C. V. Verhoosel, M. A. Scott, T. J. R. Hughes, and C. M. Landis. A phase-field description of dynamic brittle fracture. *Computer Methods in Applied Mechanics and Engineering* 217–220:77–95, 2012.
3. Boresi, A. P., and R. J. Schmidt. Advanced Mechanics of Materials. Wiley, 2003, 708 pp.
4. Campbell, J. D. On the theory of initially tensioned circular membranes subjected to uniform pressure. *The Quarterly Journal of Mechanics and Applied Mathematics* 9:84–93, 1956.
5. Gent, A. N., and L. H. Lewandowski. Blow-off pressures for adhering layers. *Journal of Applied Polymer Science* 33:1567–1577, 1987.
6. Griffith, A. A. VI. The phenomena of rupture and flow in solids. *Philosophical Transactions of the Royal Society of London. Series A, Containing Papers of a Mathematical or Physical Character* 221:163–198, 1921.
7. Gültekin, O., S. P. Hager, H. Dal, and G. A. Holzapfel. Computational modeling of progressive damage and rupture in fibrous biological tissues: application to aortic dissection. *Biomech Model Mechanobiol* 18:1607–1628, 2019.
8. Li, B., and N. Bouklas. A variational phase-field model for brittle fracture in polydisperse elastomer networks. *International Journal of Solids and Structures* 182–183:193–204, 2020.
9. Roach, M. R., J. C. He, and R. G. Kratky. Tear propagation in isolated, pressurized porcine thoracic aortas. *Can J Cardiol* 15:569–575, 1999.
10. Roccabianca, S., C. A. Figueroa, G. Tellides, and J. D. Humphrey. Quantification of regional differences in aortic stiffness in the aging human. *Journal of the Mechanical Behavior of Biomedical Materials* 29:618–634, 2014.
11. Tanné, E., T. Li, B. Bourdin, J.-J. Marigo, and C. Maurini. Crack nucleation in variational phase-field models of brittle fracture. *Journal of the Mechanics and Physics of Solids* 110:80–99, 2018.
12. The CGAL Project. CGAL User and Reference Manual. CGAL Editorial Board, 2021.at <https://doc.cgal.org/5.3/Manual/packages.html>
13. Williams, M. L. The continuum interpretation for fracture and adhesion. *Journal of Applied Polymer Science* 13:29–40, 1969.
14. Wriggers, P. Special Finite Elements for Continua. In: Nonlinear Finite Element Methods. Berlin, Heidelberg: Springer, 2008, pp. 399–460.doi:10.1007/978-3-540-71001-1_10
